# Automated evaluation of quaternary structures from protein crystals

**DOI:** 10.1101/224717

**Authors:** Spencer Bliven, Aleix Lafita, Althea Parker, Guido Capitani, Jose M Duarte

## Abstract

A correct assessment of the quaternary structure of proteins is a fundamental prerequisite to understanding their function, physico-chemical properties and mode of interaction with other proteins. Currently about 90% of structures in the Protein Data Bank are crystal structures, in which the correct quaternary structure is embedded in the crystal lattice among a number of crystal contacts. Computational methods are required to 1) classify all protein-protein contacts in crystal lattices as biologically relevant or crystal contacts and 2) provide an assessment of how the biologically relevant interfaces combine into a biological assembly In our previous work we addressed the first problem with our EPPIC (Evolutionary Protein Protein Interface Classifier) method. Here, we present our solution to the second problem with a new method that combines the interface classification results with symmetry and topology considerations. The new algorithm enumerates all possible valid assemblies within the crystal using a graph representation of the lattice and predicts the most probable biological unit based on the pairwise interface scoring. Our method achieves 85% precision on a new dataset of 1,481 biological assemblies with consensus of PDB annotations. Although almost the same precision is achieved by PISA, currently the most popular quaternary structure assignment method, we show that, due to the fundamentally different approach to the problem, the two methods are complementary and could be combined to improve biological assembly assignments. The software for the automatic assessment of protein assemblies (EPPIC version 3) has been made available through a web server at http://www.eppic-web.org.

**Author summary:** X-ray diffraction experiments are the main experimental technique to reveal the detailed atomic 3-dimensional structure of proteins. In these experiments, proteins are packed into crystals, an environment that is far away from their native solution environment. Determining which parts of the structure reflect the protein’s state in the cell rather than being artifacts of the crystal environment can be a difficult task. How the different protein subunits assemble together in solution is known as the quaternary structure. Finding the correct quaternary structure is important both to understand protein oligomerization and for the understanding of protein-protein interactions at large. Here we present a new method to automatically determine the quaternary structure of proteins given their crystal structure. We provide a theoretical basis for properties that correct protein assemblies should possess, and provide a systematic evaluation of all possible assemblies according to these properties. The method provides a guidance to the experimental structural biologist as well as to structural bioinformaticians analyzing protein structures in bulk. Assemblies are provided for all proteins in the Protein Data Bank through a public website and database that is updated weekly as new structures are released.

## Introduction

It has been known for nearly a century that many proteins are complex assemblies of polypeptide subunits [1] and the protein quaternary structure terminology was first formalized by J.D. Bernal in the late 50s [2]. Over the following decades, the importance of quaternary structure became fully appreciated, especially thanks to the transformative technological advances that led to the structure determination of more than hundred thousand proteins. In the Protein Data Bank (PDB) [3] about 50% of structures are annotated as monomeric (62887/132661, as of Aug 15, 2017). This clearly illustrates the importance of correctly interpreting and assigning quaternary structure. Fundamentally, the quaternary structure of proteins determines their physiochemical behavior and mode of interaction with other molecular partners, eventually contributing to their biological function.

In structures determined by X-ray crystallography (89% of the current PDB), the biologically relevant interfaces building the quaternary structure are embedded in a crystal lattice that contains a much larger number of non-biologically relevant crystal contacts. A recent comprehensive study of all protein-protein contacts in the PDB estimates that the ratio of biologically relevant interfaces to crystal contacts is about 1 in 6 [4]. Crystallographic techniques do not distinguish between the two kinds of contacts, and common experimental methods such as size-exclusion chromatography reveal only the stoichiometry of the complex rather than the detailed binding mode [5]. Due to this difficulty, the error rate in biological assembly annotations in the PDB has been estimated as at least 7% [4] and as much as 14% [6]. Such errors can significantly effect downstream uses of protein structures that assume the correct assignment of the biological assembly (for instance, structure prediction or docking). As such, computational tools are needed to determine biological assemblies in crystallographic structures to identify errors and better annotate novel structures.

We have recently reviewed the protein interface classification problem and the theoretical and software solutions devised over the years to address it [5]. Previous approaches have relied on structural properties (PITA [7]), thermodynamic estimation of interface stability (PISA [8]), machine learning (IPAC [9], [10]), and comparison to other proteins (PiQSi [6], ProtCID [11]). Our own method, the Evolutionary Protein-Protein Interface Classifier (EPPIC, [12]), utilizes information about the evolutionary conservation of interface residues to classify interfaces.

Previous versions of EPPIC have focused on the classification of individual interfaces; here we extend EPPIC to consider the crystal lattice as a whole and identify the biological assembly therein. Once all protein-protein interfaces in a given crystal structures have been classified, the information can be combined to infer a consistent biological assembly for the crystal. In many cases the biological unit assignment is clear cut, for instance when all interfaces are clearly classified as crystal packing. However, there are also many cases where assigning a clear biological unit is far from trivial even for the experienced structural biologist. The correct choice depends not only on a good-quality classification of the interfaces involved but also on symmetry and topology considerations. In those cases, a computational assessment is an extremely useful tool. It is also important for systematic, PDB-wide studies of protein quaternary structures, since it removes the partly subjective character of human assignments.

The new tool provides a comprehensive enumeration of all valid assemblies in a protein crystal lattice, taking into account topology and symmetry considerations, presenting an effective and comprehensive prediction of protein quaternary structures. To the best of our knowledge, our method is the first that automatically enumerates all the valid assemblies present in a protein crystal. In the following sections we present the method, its implementation and performance and explain the different issues that arise.

### The biological assembly

The PDB defines the biological assembly as “the macromolecular assembly that has either been shown to be or is believed to be the functional form of the molecule” [13]. Many proteins do have a single clear functional unit which accounts for the majority of the folded species in cells. However, determining the biological assembly in crystals can be less clear-cut. Modifications in the protein construct to facilitate crystallization, such as removal of disordered loops or domains, can alter or remove interfaces, giving a different assembly than would be present *in vivo*. In such cases, rather than representing the functional form of the molecule, the best we can hope for is representing the complex that would remain were the crystal to be dissolved in a physiological-like buffer.

Weak interactions represent a further challenge. Many protein-protein interactions can be described as transient or weak, as measured by a high dissociation constant (*K_d_*). The crystal environment may or may not capture those transient assemblies. EPPIC is targeted at predicting stable biological assemblies; in cases where the protein is likely to exist in equilibrium under physiological conditions, we consider both states to be correct biological assemblies and typically predict the smaller (more stable) assembly. However, this is not a major issue in practice, as all cases considered in the benchmark had a clear consensus as to the correct biological assembly.

### Definitions

Let us first introduce a few definitions that will be used throughout the manuscript:

1. A **molecular entity** is a unique molecule (typically a polypeptide) with an unique sequence. Different instances (chains) of the entity can have slight differences in 3D conformation, as it is often the case with non-crystallographic symmetry (NCS) copies of the same molecule.
2. An **interface type** is a particular binding mode between two entities. Since the atomic details of an interface may differ between two instances of an interface type in a particular crystal, a clustering protocol over the set of inter-chain contacts is required to define equivalent interfaces.
3. An **assembly** is a set of chains from the unit cell (lattice). Chains are identified by a chain ID and a symmetry operator ID.
4. An interface between two given chains is **engaged** in an assembly if both partners of the interface belong to the assembly.
5. An interface can be considered **induced** if when disengaging it, the assembly remains connected. It can be seen as a “redundant” interface in the assembly. A minimum of 1 interface type is needed in a cyclic point group symmetry, or 2 interface types for other point groups (dihedral, octahedral, tetrahedral, icosahedral). All other interfaces can be considered induced. The choice of what is a constitutive interface and what is an induced interface is subjective. See for instance Figure 1b where the dihedral assembly is composed of 3 interface types, with 1 of them being induced.
6. A **superassembly** is a set of assemblies that completely cover the lattice.
7. The **stoichiometry** of an assembly is a positive integer vector containing the molecule counts for each entity in that assembly. For example, in a crystal with entities *A,B,C* an *A*_2_ dimer would have stoichiometry [2,0,0].
8. Two assemblies are **orthogonal** if and only if they do not have any entities in common (i.e. the inner (scalar) product of their stoichiometries is zero).

**Fig 1.**
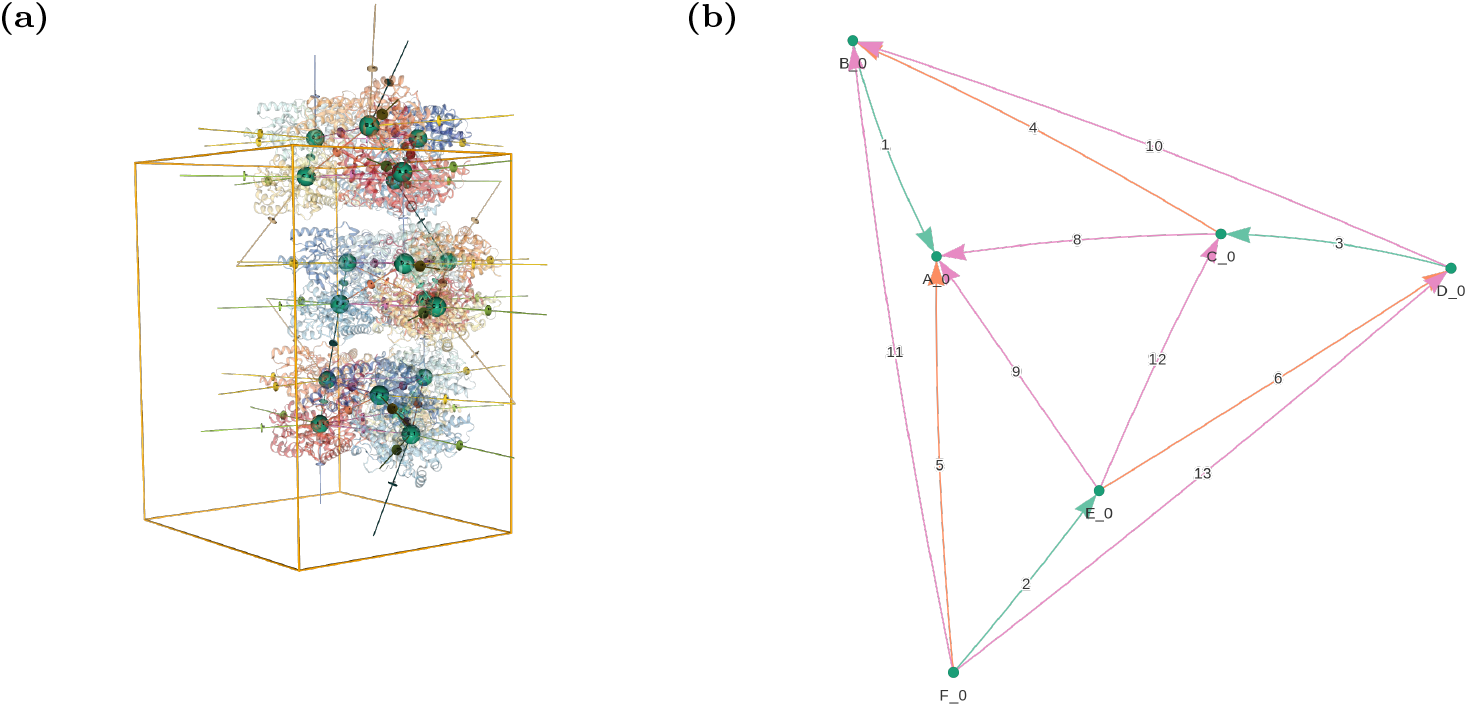
Visualizations of the biological assembly for GAD1 from Arabidopsis thaliana [PDB:3HBX] [20], as presented by the EPPIC server. (a) 3D lattice graph of a full unit cell (http://eppic-web.org/ewui/ewui/latticeGraph?id=3hbx&interfaces=*). The nodes are placed at the centroids of each chain, with edges indicating all interfaces. Many edges extend outside the unit cell due to the periodic nature of the lattice. (b) 2D graph of the hexameric biological assembly, formed by engaging three interface types (interfaces 1-3, 4-6 and 8-13). In both diagrams, nodes are labeled with chain ID and symmetry operator and colored by molecular entity. Edges are numbered sequentially by buried surface area and colored by interface type.

### Symmetry and closeness

In their seminal paper, Monod, Wyman and Changeux [14] exposed the basics of protein association into oligomers by presenting a very clear argumentation on the possible ways in which homomers can associate. They argue that only two types of associations are possible between two protein chains of the same entity:

1. **Isologous** or face-to-face: two protomers meet through a 2-fold symmetry axis. The same surface patch is used for the association.
2. **Heterologous** or face-to-back: two different surface patches from each side mediate the association.

In isologous associations the interacting interface patches are mutually satisfied and capped. There is no further association possible through the interfaces. However in heterologous association the interacting interface patches are exposed to the solvent and will continue associating to other protomers indefinitely. The only way that this indefinite association can stop is by the protomers cycling around and associating back to the first protomer, forming a cyclic *C_n_* symmetry. Thus in both cases, in order to have stable oligomeric complexes in solution, symmetry must occur. Specifically, point group symmetry is necessary: cyclic (*C*), dihedral (*D*), tetrahedral (*T*), octahedral (*O*), or icosahedral (*I*). Cyclic is the only point group that is composed by only heterologous interfaces, while the others are combinations of both isologous and heterologous interfaces.

The same argument can be extended to heteromers with two or more copies of each monomer. The heteromer is reduced to the homomer case by simply fusing the heteromeric entities into one and then treating the super entity as a homomer. Symmetry is thus a necessary condition for stable protein oligomers and we found our subsequent analysis and the assembly rules on that assumption.

The necessity and prevalence of symmetry has been since widely studied in the literature. The review by Goodsell and Olson [15] is a comprehensive overview of the topic. There are mechanisms that can lead to non-symmetric assemblies, for instance pseudo-symmetry or self-occlusion producing steric hindrance on an heterologous interaction [16]. However those exceptions are rare and the vast majority of known protein oligomers are symmetric. We discuss some of the exceptions in the section Exceptions to the rules below.

### The lattice graph

The crystal lattice can be represented by a periodic graph with protein chains as nodes and interfaces between them as edges. Graphs that represent lattices are widely used in crystallography (especially for small molecules) and are also known as crystal nets. The excellent book by Sunada [17] contains an in-depth account of the mathematics of crystal nets. Here, we apply them to whole macromolecules rather than individual atoms and bonds, as is more typical in small molecule crystallography.

We label nodes and edges to identify the molecular entities and the distinct mode of interactions between them, see Figure 1b. A node is identified by a chain identifier and a symmetry operator identifier (e.g. A_1), while an edge is identified by a numerical interface identifier. Additionally all nodes corresponding to the same molecular entity are given an entity label and all edges corresponding to the same interface type are given an interface type identifier label. Although the graph is depicted in one unit cell only, it does represent all possible connections in the crystal including those across neighboring unit cells.

The crystal translations associated to the interfaces are also required to fully describe the graph and are essential in finding closed cycles with 0 net translation. We represent these as an integer vector for each edge giving the difference in Miller indices for the two chains participating in the interface, with respect to a given choice of unit cell operators.

Diagrams similar to our 2D graph representation of the lattice graph have been used previously in the context of quaternary structure studies, see for instance [18] and [19]. However in those studies the diagrams lacked information on interface types and their connectivity, for instance sometimes representing a D3 assembly with only one kind of edge.

## Methods

### Assembly rules

Given the definitions introduced in the section above, we now establish the rules for a superassembly to be valid, from which the algorithm to find all assemblies result:

1. **Full Coverage:** Every chain belongs to exactly one assembly from the superassembly (implied from the definition).
2. **Uniform Composition:** All instances of an interface type are either engaged or disengaged in the superassembly.
3. **Isomorphism:** All assemblies should have isomorphic graphs with respect to the molecular entities and interface types. Isomorphism must hold only if the assemblies are not orthogonal in stoichiometry.
4. **Closed Symmetry:** No combination of engaged interface operators can lead to a non-zero pure translational operator.

The first two rules ensure consistency in the decomposition of the superassembly into assemblies. The third rule is motivated by the assumption that cocrystallization of multiple biological assemblies involving the same entities does not occur. Co-crystallization implies that the complex exists at equilibrium in the crystallization conditions, making the correct biological assembly ambiguous. By disallowing co-crystallization we effectively favor the dissociated form as the correct assembly for proteins with weak or transient interactions. Finally, the fourth rule is motivated by the hypothesis that infinite assemblies are never biological (discussed later).

From the rules it follows that a) valid assemblies are point group symmetric, and b) heteromeric assemblies must have even stoichiometry. We then implement an algorithm that follows the above rules, described in detail below.

### Pairwise interface classification

Interface classification in EPPIC is described in our previous paper [12]. However, there have been some improvements to the interface scoring and classification.

When calculating the sequence entropy at each position, we now use a 6-letter reduced alphabet to represent the 20 amino acids [21]. The alphabet was proposed by Mirny et al. [22]:

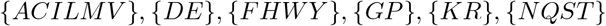

In addition, the core surface scores are now pure Z-scores where *m* residues are sampled 10,000 times from the whole protein surface. An average sequence entropy is calculated for each of those samples and then the mean and standard deviation of the whole distribution is used for the Z-score of the *m* residues composing the interface core.

Finally, we have introduced a probabilistic scoring for interface classification, based on a logistic regression classifier that uses 2 of our 3 previous indicators: geometry (gm) and core-surface (cs) scores [23]. The model was trained using the *Many* dataset [4] with R generalized linear model (glm) functions. The equation that describes the probability of an interface being biologically relevant (p) is:

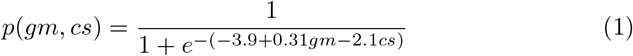

A ROC curve with the performance of the new method can be found in Supplementary Figure 1, directly comparable to the curves in [4].

### Assembly enumeration algorithm

We denote interface types by numerical identifiers 1,…, *n*, sorted from largest to smallest area. An assembly is created when engaging a subset of those interfaces, e.g. {1, 3} is the assembly where only interfaces 1 and 3 are engaged, or {} is the empty assembly where no interfaces are engaged.

Given the set of all interface types *S* = {1,…, *n*}, enumerating all possible assemblies is a matter of traversing the tree of its power set 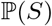. A total of 2^*n*^ assemblies are possible in principle, making the full enumeration prohibitive when *n* becomes large. For every set, the assembly is tested against our rules to see if it represents a valid assembly. An important observation makes the problem more tractable: if a given set is invalid, all of its children (i.e. any other set that contains the same engaged interfaces plus any other) will also be invalid. This dramatically prunes the tree, making it possible to quickly do the exhaustive enumeration for almost all cases.

As a further optimization, heteromers with many protein entities are reduced to equivalent homomeric lattice graphs by combining entities, leading to considerably simpler graphs. Interfaces that join different entities are selected in a greedy manner. The edge corresponding to the interface is then contracted, merging the two entities into a single node. This process is iterated until a single meta-entity remains. Graph contraction preserves the structure of the graph with respect to the validity properties and relative score, while allowing considerably faster superassembly enumeration.

The test of validity for a given superassembly boils down to two tests: graph isomorphism and finding closed cycles in the graph. To find the cycles we use the Paton algorithm [24] as implemented in the JGraphT library.

The EPPIC software package implements all of the described algorithms in its new version 3. The software is written in Java, using BioJava [25] as the underlying software library to handle the biological data.

### Predicting assemblies from pairwise interface classification

In order to predict the most likely biologically relevant assembly we use a combination of the probabilistic scores calculated for the pairwise interfaces. We consider an interface as a binary event, that can either occur or not in biological conditions. An assembly is just a subset of the interfaces in the crystal occurring in biological conditions, with the remaining interface subsets not occurring. If we denote by *S* the set of all protein interfaces, each assembly can be defined as a boolean vector of length |*S*|, with *s_i_* indicating that the interface *i* is engaged in the assembly. The probability *p_i_* of an interface *i* being biologically relevant is calculated per equation 1. To estimate the probability of an assembly occurring in biological conditions is to estimate the joint probability of events coming from all interfaces in the crystal:

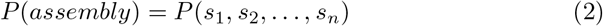

To perform the estimation, we assume that the pairwise interfaces can be treated as independent. As EPPIC interface scores depend critically on the estimation of residue burial, this assumption is valid for the score in equation 1 so long as the total buried surface area of the assembly is well approximated by the sum of the pairwise buried surface areas.

Using the probabilities for each interface *p*_*i*_, we can assign a probability of occurring in biological conditions to each assembly of the crystal:

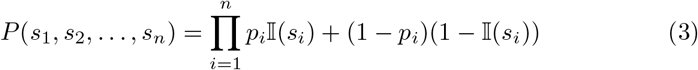

where 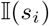 is the indicator function 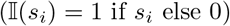.

Some combinations of engaged interfaces will correspond to invalid assemblies according to the rules above. These assemblies have a probability of occurrence of 0, so summing *P*(*S*) over all valid assemblies in the crystal may be less than one. Thus, a final normalization step can be applied to redistribute the probability mass of interface events leading to invalid assemblies into the valid assemblies.

Special care has to be taken with induced interfaces, which can be omitted from an assembly without changing the quaternary structure. Superassemblies which differ only by an induced interface can be easily detected by comparing the stoichiometry of their constituent assemblies. This allows all superassemblies which differ only by induced interfaces to be combined together. The superassembly with the highest number of engaged interfaces is reported along with the total probability of all equivalent superassemblies.

The reported probability for an assembly is the confidence that the EPPIC call is correct. It is important not to confuse these probabilities with strength of the assembly or transitivity properties.

### Biological assemblies dataset

We compiled a new dataset of biological assemblies using the annotations of deposited structures in the PDB. We started with 96,594 crystal structures with higher than 3 Å resolution and lower than 0.3 R-free value from the PDB. Structures were then grouped into 60,034 unique sequences and 36,843 70% sequence identity clusters for each of their chains. These were further filtered to clusters with at least three structures and where all structures had the same biological assembly annotation. Randomly selecting a representative from each of the remaining clusters yielded 1,481 proteins. This new dataset represents a diverse sample of the PDB: 53% of oligomers, from which 11% are heteromers, covering macromolecular sizes up to 24 partner subunits.

### Improvements in the web server

Together with the command line interface (downloadable at http://eppic-web.org/ewui/ewui/latticeGraph?id=3hbx&interfaces=*), we provide a web server with a graphical user interface to the EPPIC 3 software. There has been numerous improvements compared to what we described earlier.

A new view provides the full enumeration of all valid assemblies found in the crystal structure with links to its constituent interfaces. The assemblies are visualized by thumbnail images of the assembled proteins and by 2-dimensional diagrams of their corresponding graphs.

New lattice graph visualizations are provided. First in 2D with the help of the vis.js library [26]. An optimal 2D graph layout is achieved by performing a stereographic projection of the 3D molecule. A 3D lattice graph representation is also provided with NGL [27] by overlaying custom made spheres and cylinder objects on top of a semi-transparent cartoon representation of the unit cell.

In EPPIC 2, the 3D visualization was based in the Jmol molecular viewer. The server now uses NGL [27] as the molecular visualization software. NGL is written in JavaScript and runs natively in the browser with very good performance thanks to WebGL technologies. Its advanced features allow for showing sequence entropy surface color representation within the browser.

## Results and Discussion

Our approach to find quaternary structure assemblies in a given crystal structure is primarily based in the representation of the lattice as a periodic graph. The fundamental assumption is that an assembly needs to exhibit point group symmetry in order to be valid. The point group symmetry requirement is equivalent to finding certain closed paths in the lattice graph, as described in the Assembly rules section above. The algorithm is thus able to enumerate all topologically valid assemblies in the crystal. Subsequently scoring the different viable assemblies is based on a combination of the pairwise interface scores.

### Benchmarking and comparison with PISA

We validated the assembly assignment method against the 1,481 PDB entries with consensus quaternary structure annotations. Figure 2 shows the confusion matrix of the assembly size for EPPIC predictions, with an overall precision of 85%. While the precision is constant across the different macromolecular sizes, the recall is lower for larger assemblies. The consequence is the reduction of non-biological large macromolecular assembly predictions (top-left of the matrix in Figure 2), at the expense of predicting some partial assemblies (bottom-right of the matrix in Figure 2).

**Fig 2.**
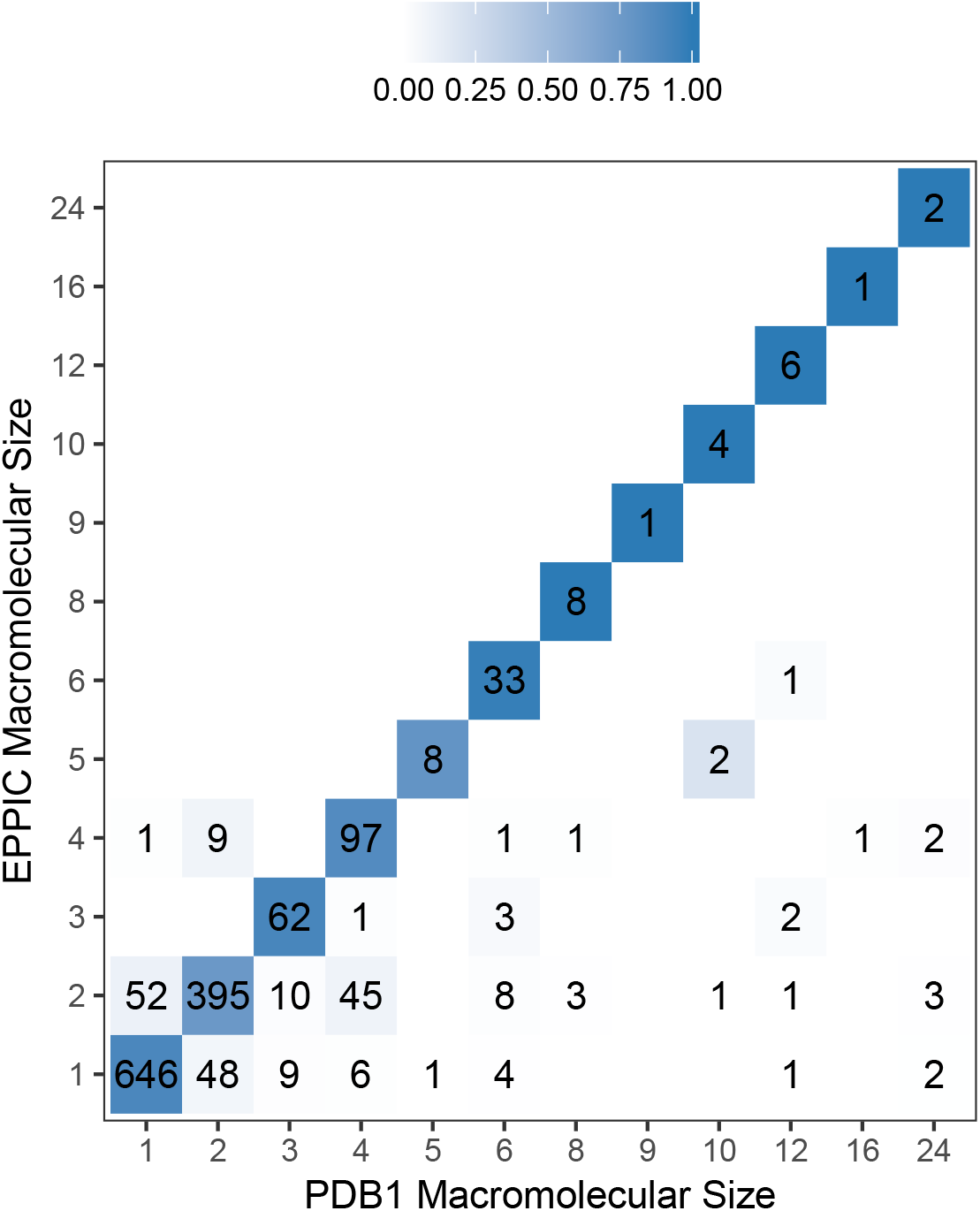
EPPIC assembly predictions as a confusion matrix of macromolecular sizes. Tiles are colored as the fraction of predictions (i.e. row normalized). The method achieves 85% precision on the dataset. PDB1 refers to the 1st biological assembly annotation provided by the PDB, in here considered as the true biological assembly.

As a further validation, we provide a comparison to the popular PISA method, the de-facto standard in the field. Despite very similar overall precision in the assemblies dataset, EPPIC and PISA predictions show many differences, as it can be appreciated in Figure 3. The most important difference is that PISA makes the opposite trade-off in the prediction of large macromolecular size assemblies, achieving better accuracy for larger assemblies at the expense of predicting some non-biological large assemblies. Table 1 gives the overview of over and under predictions, whilst Table 2 contains more detailed statistics divided into 3 categories: monomers, dimers and higher oligomers.

**Table 1.**
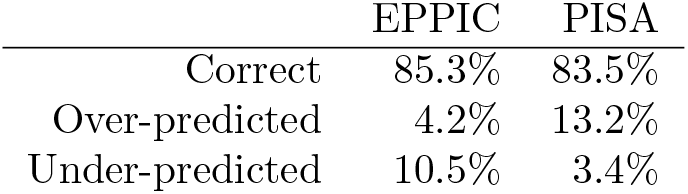
Over and under predictions as a summary of Figure 3. Over predictions correspond to the upper-left half of the confusion matrix whilst under predictions correspond to the lower-right half.

|  | EPPIC | PISA |
| --- | --- | --- |
| Correct | 85.3% | 83.5% |
| Over-predicted | 4.2% | 13.2% |
| Under-predicted | 10.5% | 3.4% |

**Table 2.** Prediction statistics for different categories: monomers, dimers and higher-oligomers (macromolecular size ≥ 3).

|  | Precision |  | Recall |  |
| --- | --- | --- | --- | --- |
|  | EPPIC | PISA | EPPIC | PISA |
| Monomers | 90% | 96% | 92% | 82% |
| Dimers | 76% | 75% | 87% | 84% |
| Higher Oligomers | 90% | 76% | 67% | 86% |

**Fig 3.**
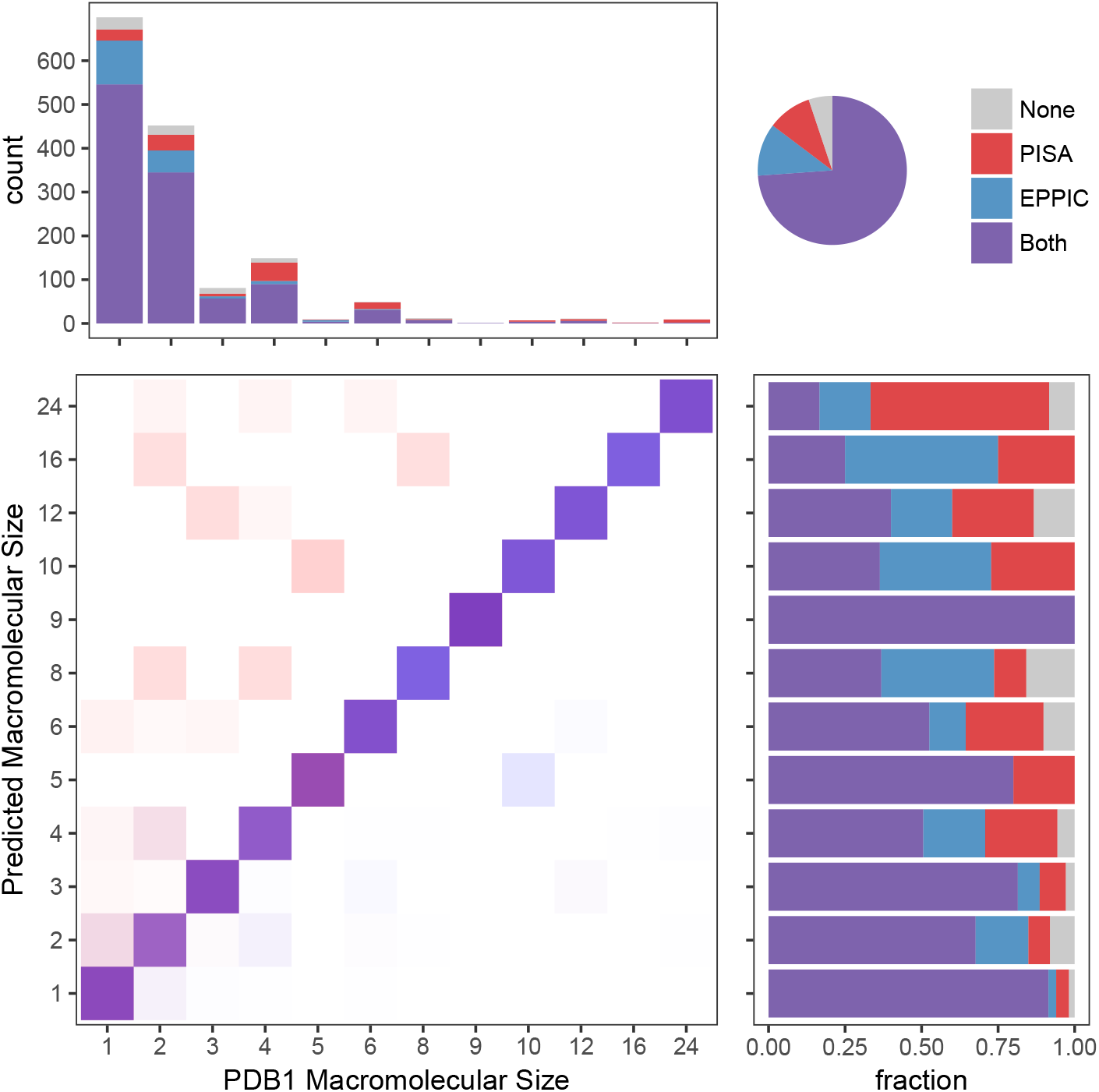
Comparison of assembly predictions from EPPIC and PISA on the benchmarking dataset. On the top right, a pie chart shows the global agreement between EPPIC and PISA. On the bottom left, the confusion matrix of actual (PDB1 annotations) and predicted macromolecular sizes. Tiles colored as a fraction of each EPPIC (blue) and PISA (red) macromolecular size prediction (i.e. row normalized). On the bottom right, the agreement and precision of the methods for each PISA macromolecular size prediction. On the top left, the total number and recall for each macromolecular size in the dataset.

The agreement of the two methods greatly increases the confidence of a prediction. As observed in Figure 4, when EPPIC and PISA agree, in 78% of the cases, the error rate is only 5%. On the other hand, when the methods disagree, in the remaining 22% of the structures, the error rate of each method is around 50%. Therefore, each method corrects roughly the same amount of assignments of the other. Furthermore, at least one of the two methods is correct in 95% of the cases. These results suggest that a meta-method combining EPPIC and PISA could be successful, with a potential precision of up to 95%.

**Fig 4.**
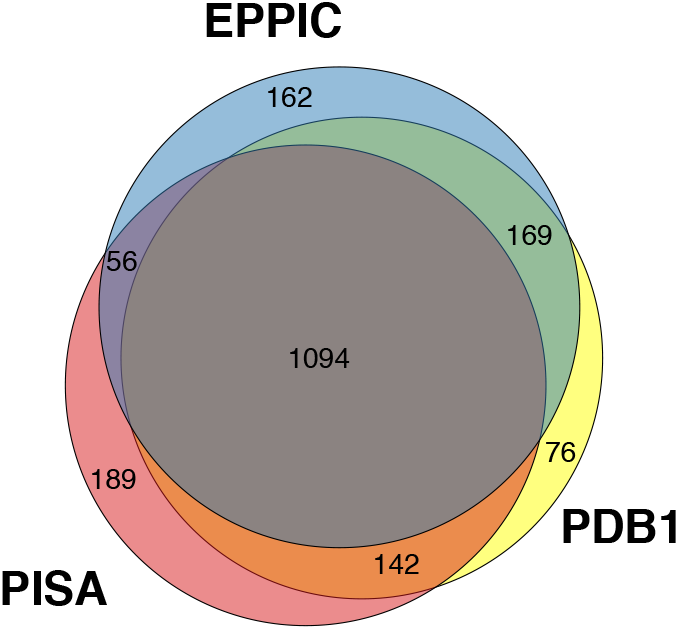
EPPIC and PISA predictions on the protein assembly dataset as a Venn diagram. PDB1 refers to the 1st biological assembly annotation provided by the PDB.

Additionally to the benchmark with our dataset we have also measured the performance with the PiQSi dataset [6], composed of 1315 biological assemblies curated with a combination of manual community annotation and automatic methods. The precision values for the PiQSi benchmark are 73% for EPPIC and 79% for PISA. It should be noted that the PiQSi dataset is less representative of the PDB compared to our dataset, for instance having fewer monomers and more very large oligomers than what is average in the PDB.

### Interesting assemblies in the PDB

In most cases, the quaternary structure interpretation of a crystal is unambiguous to a trained crystallographer. The unit cell shows clear blocks of symmetrically packed molecular entities. However, in more difficult cases the interpretation of the crystal is far from obvious and requires very careful observation.

A good example is the crystal structure of the fimbrial adhesin FimH protein (PDB 2VCO [28]). The crystal contains two FimH molecules in the asymmetric unit interacting via a heterologous interface. All other interfaces in the crystal are also heterologous, except for the very weak isologous interface 6 (as identified by EPPIC, see http://eppic-web.Org/ewui/#interfaces/2vco). No combination of the interfaces produces a closed cycle (assembly rule 4 is not satisfied). Thus the only valid assembly in the crystal is monomeric (see http://eppic-web.org/ewui/#id/2vco). However, the PDB annotation for this case engages interfaces 1 and 3 to form a tetramer. The global symmetry of the tetramer, as calculated by the RCSB PDB website [29], is *C*_2_, indicating that the tetramer is not point group symmetric (the only possible point groups for an *A*_4_ stoichiometry are *C*_4_ or *D*_2_). The assembly might seem reasonable since in the crystal it shows as an independent block repeated throughout (see Figure 5a). Figure 5b helps explain this with a simple 2D schematic representation of a crystal packing with heterologous interfaces. The PISA software predicts in this case a different tetrameric assembly than the one annotated in PDB, formed by engaging interfaces 1 and 6. Again this assembly does not contain point group symmetry. This example also shows how a simple search for stoichiometry-symmetry imbalance (i.e. *A_n_* stoichiometry should have *C_n_* or *D_n_*/_2_ point group symmetry) would uncover similar cases of potentially erroneous annotations in the PDB.

**Fig 5.**
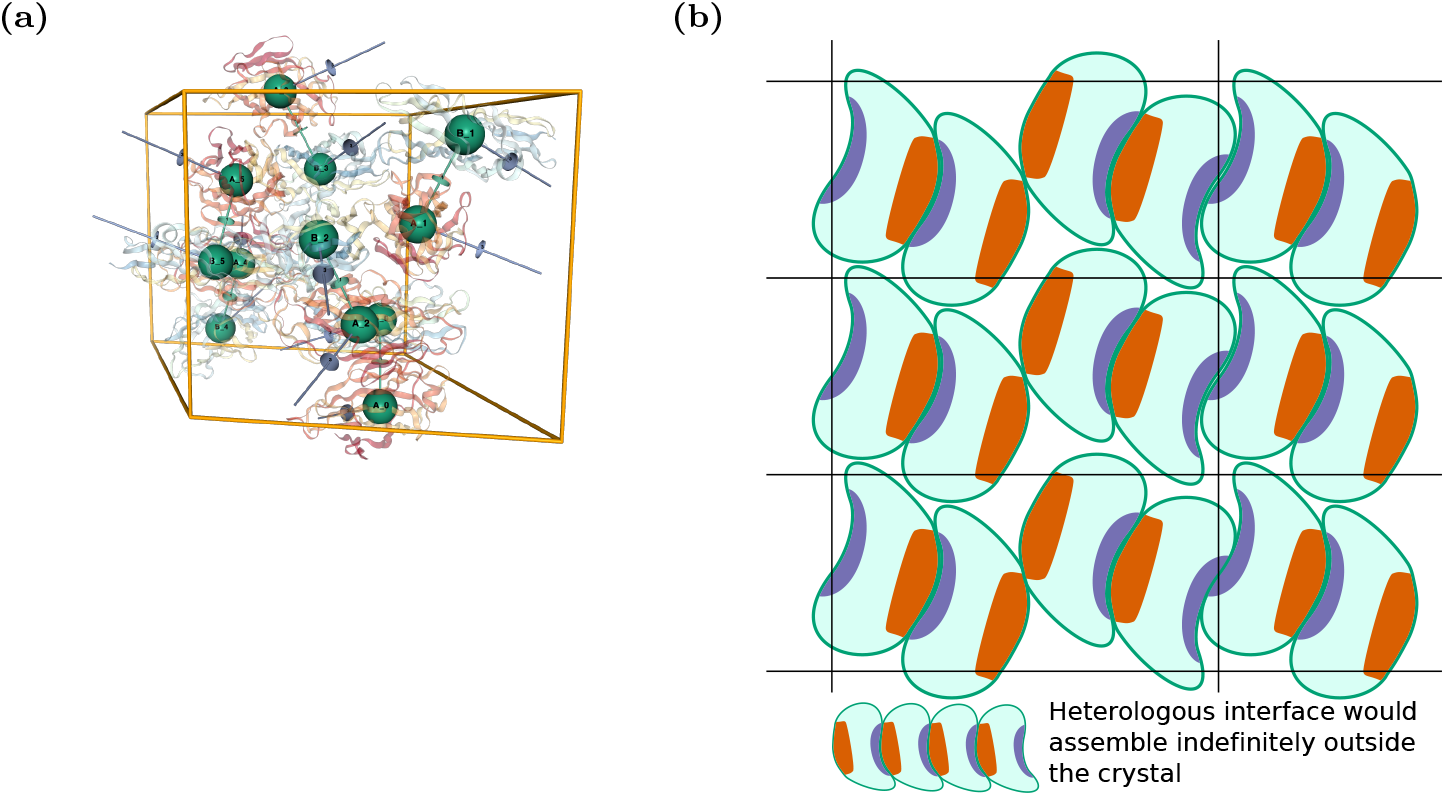
(a) The crystal lattice of PDB 2VCO as shown by the EPPIC server (http://eppic-web.org/ewui/ewui/latticeGraph?id=2vco&interfaces=1,3). The highlighted tetrameric assembly is the one annotated in the PDB. (b) Schematic 2D representation of a lattice that contains an asymmetric dimer through a heterologous interface but which does not form infinite fibers in the crystal.

Another similar example is lipoteichoic acid synthase LtaP from Listeria monocytogenes (PDB 4UOP), which corresponds quite closely to the schematic representation of Figure 5b: 2 molecules in the asymmetric unit interact through a heterologous interface, with the heterologous interface capped in the crystal by other molecules. The PDB annotates the asymmetric dimer in the AU as the biological assembly based on a PISA prediction. However, the protein is known to be a monomer in solution based on size exclusion chromatography [30]. Since the dimeric assembly is not symmetric, EPPIC considers it invalid following the assembly rules.

A second example of a subtle lattice that is difficult to analyze manually would be that of the crystal structure of the putrescine receptor PotF from E. Coli (PDB 1A99 [31]; see Figure 6a). There are 4 PotF molecules in the asymmetric unit. Two different isologous interfaces relate the 4 molecules in the AU, interfaces 5 (D+C) and 6 (B+A). The PDB annotates a dimeric assembly through one of the interfaces in the asymmetric unit (interface id 6). In principle, the assembly is valid since it has *C*_2_ point group symmetry. However, a more careful analysis of the crystal shows that not all monomers in the lattice participate in this kind of interaction: the C and D chains do not interact in the same way throughout the crystal. Considering this assembly as a dimer would break the full coverage rule (rule 1), while considering it a co-crystal dimer + monomer breaks the isomorphism rule (rule 3). This shows why isomorphism is important: a stable assembly in solution can not occur only in some parts of the lattice and not in others. The schematic 2D view of Figure 6b helps visualize the problem. By following the assembly rule, EPPIC finds here only a monomeric assembly (see http://eppic-web.org/ewui/#id/1a99). In this case PISA predicts a disjoint assembly formed by a *A*_2_*B*_2_ tetramer and separate monomers of chains C and D.

**Fig 6.**
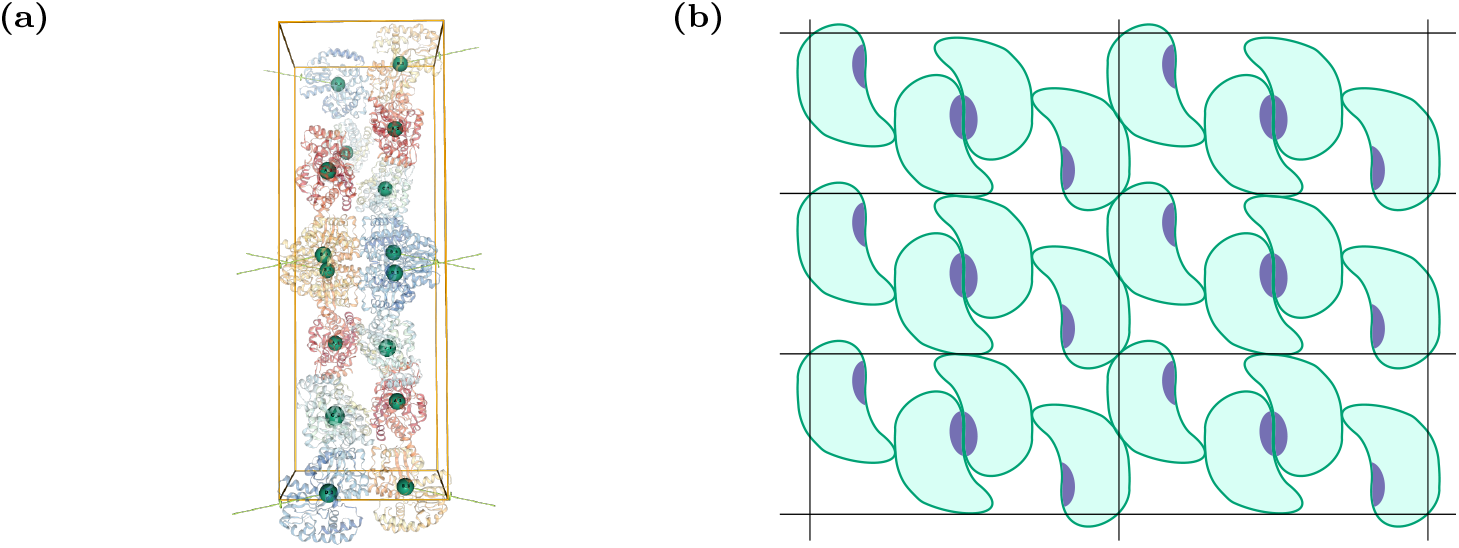
(a) The crystal lattice of PDB 1A99, highlighting the C_2_ dimer wrapping around the unit cell (http://eppic-web.org/ewui/ewui/latticeGraph?id=1a99&interfaces=7). (b) Schematic 2D representation of a lattice that contains a valid C_2_ assembly, but which is not isomorphic throughout the crystal.

### Exceptions to the rules

Non-symmetric assemblies are very rare but still a possibility. In fact as of June 2017, 96% of PDB structures are annotated with symmetric biological assemblies. A comprehensive study of asymmetric assemblies in heteromers [16] found a similar fraction of asymmetric cases for heteromers (9.8% of all heteromers have uneven stoichiometry). In their in-depth study, a thorough review of all cases unearthed a number of quaternary structure assignment errors, further lowering the asymmetric fraction.

Different mechanisms can lead to breakage of symmetry. One major cause of exceptions is the existence of pseudosymmetry in heteromers with uneven stoichiometry (e.g. PDB 4FI3 [32]), whereby one entity can bind several copies of its partner at distinct but structurally similar binding sites. Other exceptions include steric hindrance (e.g. PDB 3Q66 [33]) and extreme conformational flexibility (e.g. PDB 1YGY [34]). An additional source of exceptions is filamentous proteins and amyloids, which violate rule 4 by definition. However, since these properties make them resistant to crystallization, such cases are rare.

A prominent example of a pseudosymmetric case is that of the B12 vitamin transporter [32]. This large membrane protein complex is composed of 5 subunits, with 3 distinct molecular components (Figure 7a). Two BtuD chains form a symmetric *C*_2_ dimer in the cytoplasmic domain, while the transmembrane domain is composed of two BtuC chains arranged along the same *C*_2_ axis. Capping the complex on the periplasmic side is a single BtuF chain that binds to the BtuC dimer in a symmetric way. The 1:2 symmetric binding is made possible by the internal pseudosymmetry of the BtuF chain (see Figure 7b and c).

**Fig 7.**
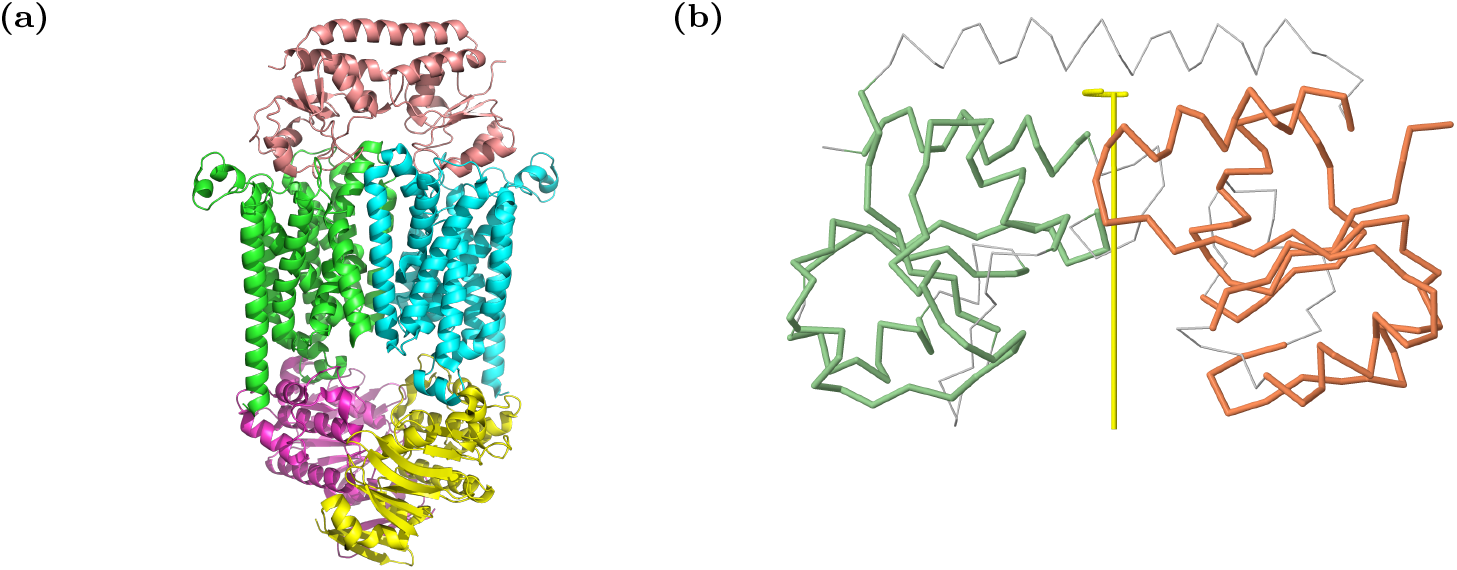
(a) The ABC transporter (PDB 4FI3). (b) The BtuF periplasmic domain with internal *C_2_* pseudo-symmetry highlighted, including the 2-fold axis of symmetry. The internal symmetry calculation was performed with CE-Symm [35]

## Conclusions

We have presented an approach to enumerate and predict quaternary assemblies from protein crystal structures. This new method should prove very useful to the crystallographer, considerably easing the assembly interpretation of protein crystals. The automated exhaustive enumeration of assemblies represents a great improvement in the quaternary interpretation of structures, which to a large extent still requires human subjective interpretation.

Our ideas are centered in the necessity of symmetry based on the simple arguments established by Monod, Wyman and Changeux [14]. Symmetry is essential for stable soluble proteins. Our method can thus help in avoiding mistaken asymmetric interpretation of assemblies. It can also serve as a validation tool for atomic models that lack symmetric or isomorphic assemblies, providing hints on possibly uninterpreted regions of electron density that need to be added to the model in order to complete it. Additionally existing methods to predict quaternary assemblies [8] are not always strict in the symmetry constraint, providing sometimes misleading interpretations of the crystal.

Importantly, our assembly scoring uses evolution as the ultimate arbiters to the biological relevance of the assemblies, making this method complementary to existing methods based on thermodynamic estimations. Also, the newly introduced confidence values provide a clear guide to interpreting the predictions. At the same time, confidence estimations provide a means to more reliably estimate biological assembly annotation errors in the Protein Data Bank, as well as aiding the crystallographer in deciding when additional oligomeric experimental evidence for a particular assembly might be needed. Confidence values also allow for fully automated analyses of oligomeric complexes at the PDB wide level.

Some new avenues of research are possible based on this new resource. For instance, the assembly graphs allow for more detailed study of different crystal lattices and their relationships across the PDB.

We also recognize that our strict enforcement of point group symmetry is not always ideal, since, as shown in the Results section, exceptions to symmetry do occur. In future work we plan to address the problem by relaxing some of the conditions in cases where interface scoring indicate an invalid assembly could be biological.

Recent publications indicate that the evolutionary approach to protein assembly prediction and classification can be significantly improved in the future. Two research lines are promising: co-evolution of inter-subunit residues in protein-protein interactions [36,37] and the evolutionary constraints of highly symmetric assemblies to avoid supramolecular assembly formation [38]. We believe that these additional sources of information can improve the performance of the classifier and confidence estimates, as we continue to advance the method.

## Acknowledgments

G.C. initiated and led the research, contributing to the initial versions of the manuscript. Tragically he sickened and passed away on the 2nd of May 2017. We are grateful to the RCSB PDB for supporting the continuation of the project, including hosting of the EPPIC service. Financial support to G.C. from the Swiss National Science Foundation (grant 31003A_140879) and the Research Committee of the Paul Scherrer Institute (grants FK-05.08.1, FK-04.09.1) is gratefully acknowledged, as is IT support from SyBIT/SIS. This research was supported in part by the Intramural Research Program of the National Center for Biotechnology Information, National Library of Medicine, National Institutes of Health (support to S.B.). Support for J.D. within the RCSB PDB comes from the National Science Foundation, the National Institutes of Health, and the Department of Energy (NSF DBI-1338415; Principal Investigator: Stephen K. Burley).

## References

1. Svedberg T. Mass and Size of Protein Molecules; 1929. Available from: http://www.nature.com/doifinder/10.1038/123871a0.

2. Bernal JD. General introduction structure arrangements of macromolecules. Discussions of the Faraday Society. 1958;25:7. doi:10.1039/df9582500007.

3. Berman HM, Westbrook J, Feng Z, Gilliland G, Bhat TN, Weissig H, et al. The Protein Data Bank. Nucleic Acids Research. 2000;28(1):235–242. doi:10.1093/nar/28.1.235.

4. Baskaran K, Duarte JM, Biyani N, Bliven S, Capitani G. A PDB-wide, evolution-based assessment of protein-protein interfaces. BMC Structural Biology. 2014;14:1–11.

5. Capitani G, Duarte JM, Baskaran K, Bliven S, Somody JC. Understanding the fabric of protein crystals: Computational classification of biological interfaces and crystal contacts. Bioinformatics. 2015;32(4):481–489. doi:10.1093/bioinformatics/btv622.

6. Levy ED. PiQSi: Protein Quaternary Structure Investigation. Structure. 2007;15(11):1364–1367. doi:10.1016/j.str.2007.09.019.

7. Ponstingl H, Kabir T, Thornton JM. Automatic inference of protein quaternary structure from crystals. J Appl Cryst. 2003;36:1116–1122.

8. Krissinel E, Henrick K. Inference of Macromolecular Assemblies from Crystalline State. Journal of Molecular Biology. 2007;372(3):774–797. doi:10.1016/j.jmb.2007.05.022.

9. Mitra P, Pal D. Combining Bayes classification and point group symmetry under Boolean framework for enhanced protein quaternary structure inference. Structure. 2011;19(3):304–312. doi:10.1016/j.str.2011.01.009.

10. Luo J, Guo Y, Fu Y, Wang Y, Li W, Li M. Effective discrimination between biologically relevant contacts and crystal packing contacts using new determinants. Proteins: Structure, Function, and Bioinformatics. 2014;82(11):3090–3100.

11. Xu Q, Dunbrack RL. The protein common interface database (ProtCID)-a comprehensive database of interactions of homologous proteins in multiple crystal forms. Nucleic Acids Res. 2011;39(Database issue):D761–70.

12. Duarte JM, Srebniak A, Scharer MA, Capitani G. Protein interface classification by evolutionary analysis. BMC bioinformatics. 2012;13(1):334. doi:10.1186/1471-2105-13-334.

13. Introduction to Biological Assemblies and the PDB Archive, RCSB Protein Data Bank; 2017. http://pdb101.rcsb.org/learn/guide-to-understanding-pdb-data/biological-assemblies.

14. Monod J, Wyman J, Changeux JP. On the nature of allosteric transitions: A plausible model. J Mol Biol. 1965;12(1):88–118. doi:10.1016/S0022-2836(65)80285-6.

15. Goodsell DS, Olson AJ. Structural symmetry and protein function. Annual review of biophysics and biomolecular structure. 2000;29:105–153. doi:10.1146/annurev.biophys.29.1.105.

16. Marsh JA, Rees HA, Ahnert SE, Teichmann SA. Structural and evolutionary versatility in protein complexes with uneven stoichiometry. Nature Commun. 2015;6:63–94. doi:10.1038/ncomms7394.

17. Sunada T. Topological Crystallography. vol. 6 of Surveys and Tutorials in the Applied Mathematical Sciences. Springer; 2013.

18. Levy ED, Pereira-Leal JB, Chothia C, Teichmann SA. 3D complex: a structural classification of protein complexes. PLoS computational biology. 2006;2(11):e155. doi:10.1371/journal.pcbi.0020155.

19. Ahnert SE, Marsh JA, Hernandez H, Robinson CV, Teichmann SA. Principles of assembly reveal a periodic table of protein complexes. Science (New York, NY). 2015;350(6266):aaa2245. doi:10.1126/science.aaa2245.

20. Gut H, Dominici P, Pilati S, Astegno A, Petoukhov MV, Svergun DI, et al. A common structural basis for pH- and calmodulin-mediated regulation in plant glutamate decarboxylase. Journal of molecular biology. 2009;392(2):334–351. doi:10.1016/j.jmb.2009.06.080.

21. Somody JC. Exploring the role of amino-acid alphabets in protein-interface classification. ETH Zürich; 2015.

22. Mirny LA, Shakhnovich EI. Universally conserved positions in protein folds: reading evolutionary signals about stability, folding kinetics and function. Journal of molecular biology. 1999;291(1):177–196. doi:10.1006/jmbi.1999.2911.

23. Lafita A. Assessment of protein assembly prediction in CASP12 & Conformational dynamics of integrin α-I domains. ETH Zürich; 2017. Available from: http://e-collection.library.ethz.ch/view/eth:50644.

24. Paton K. An Algorithm for Finding a Fundamental Set of Cycles of a Graph. Commun ACM. 1969;12(9):514–518. doi:10.1145/363219.363232.

25. Prlić A, Yates A, Bliven SE, Rose PW, Jacobsen J, Troshin PV, et al. BioJava: an open-source framework for bioinformatics in 2012. Bioinformatics (Oxford, England). 2012;28(20):2693–2695.

26. Vis.js JavaScript library; 2017. http://visjs.org/.

27. Rose AS, Hildebrand PW. NGL Viewer: a web application for molecular visualization. Nucleic acids research. 2015;43(W1):W576–W579. doi:10.1093/nar/gkv402.

28. Wellens A, Garofalo C, Nguyen H, Van Gerven N, Slattegard R, Her-nalsteens JP, et al. Intervening with urinary tract infections using anti-adhesives based on the crystal structure of the FimH-oligomannose-3 complex. PloS one. 2008;3(4):e2040. doi:10.1371/journal.pone.0002040.

29. Rose PW, Prlic A, Bi C, Bluhm WF, Christie CH, Dutta S, et al. The RCSB Protein Data Bank: views of structural biology for basic and applied research and education. Nucleic acids research. 2015;43(Database issue):D345–D356. doi:10.1093/nar/gku1214.

30. Campeotto I, Percy MG, MacDonald JT, Forster A, Freemont PS, Gründling A. Structural and mechanistic insight into the Listeria monocytogenes two-enzyme lipoteichoic acid synthesis system. The Journal of biological chemistry. 2014;289(41):28054–28069. doi:10.1074/jbc.M114.590570.

31. Vassylyev DG, Tomitori H, Kashiwagi K, Morikawa K, Igarashi K. Crystal structure and mutational analysis of the Escherichia coli putrescine receptor. Structural basis for substrate specificity. The Journal of biological chemistry. 1998;273(28):17604–17609.

32. Korkhov VM, Mireku SA, Locher KP. Structure of AMP-PNP-bound vitamin B12 transporter BtuCD-F. Nature. 2012;490(7420):367–372. doi:10.1038/nature11442.

33. Su D, Hu Q, Zhou H, Thompson JR, Xu RM, Zhang Z, et al. Structure and histone binding properties of the Vps75-Rtt109 chaperone-lysine acetyltransferase complex. J Biol Chem. 2011;286(18):15625–15629.

34. Dey S, Grant GA, Sacchettini JC. Crystal structure of Mycobacterium tuberculosis D-3-phosphoglycerate dehydrogenase: extreme asymmetry in a tetramer of identical subunits. The Journal of biological chemistry. 2005;280(15):14892–14899. doi:10.1074/jbc.M414489200.

35. Myers-Turnbull D, Bliven SE, Rose PW, Aziz ZK, Youkharibache P, Bourne PE, et al. Systematic detection of internal symmetry in proteins using CE-Symm. Journal of molecular biology. 2014;426(11):2255–2268. doi:10.1016/j.jmb.2014.03.010.

36. Ovchinnikov S, Kamisetty H, Baker D. Robust and accurate prediction of residue-residue interactions across protein interfaces using evolutionary information. eLife. 2014;3:e02030. doi:10.7554/eLife.02030.

37. Bitbol AF, Dwyer RS, Colwell LJ, Wingreen NS. Inferring interaction partners from protein sequences. Proceedings of the National Academy of Sciences of the United States of America. 2016;113(43):12180–12185. doi:10.1073/pnas.1606762113.

38. Garcia-Seisdedos H, Empereur-Mot C, Elad N, Levy ED. Proteins evolve on the edge of supramolecular self-assembly. Nature. 2017;548(7666):244–247. doi:10.1038/nature23320.

